# Validation of a murine proteome-wide phage display library for the identification of autoantibody specificities

**DOI:** 10.1101/2023.04.07.535899

**Authors:** Elze Rackaityte, Irina Proekt, Haleigh S. Miller, Akshaya Ramesh, Jeremy F. Brooks, Andrew F. Kung, Caleigh Mandel-Brehm, David Yu, Colin Zamecnik, Rebecca Bair, Sara E. Vazquez, Sara Sunshine, Clare L. Abram, Clifford A. Lowell, Gabrielle Rizzuto, Michael R. Wilson, Julie Zikherman, Mark S. Anderson, Joseph L. DeRisi

## Abstract

Autoimmunity is characterized by loss of tolerance to tissue-specific as well as systemic antigens, resulting in complex autoantibody landscapes. Here, we introduce and extensively validate the performance characteristics of a murine proteome-wide library for phage display immunoprecipitation and sequencing (PhIP-seq), to profile mouse autoantibodies. This system and library were validated using seven genetic mouse models across a spectrum of autoreactivity. Mice deficient in antibody production (*Rag2*^-/-^ and μMT) were used to model non-specific peptide enrichments, while cross-reactivity was evaluated using anti-ovalbumin B cell receptor (BCR)-restricted OB1 mice as a proof of principle. The PhIP-seq approach was then utilized to interrogate three distinct autoimmune disease models. First, serum from *Lyn*^-/-^ *IgD*^+/-^ mice with lupus-like disease was used to identify nuclear and apoptotic bleb reactivities, lending support to the hypothesis that apoptosis is a shared origin of these antigens. Second, serum from non-obese diabetic (NOD) mice, a polygenic model of pancreas-specific autoimmunity, enriched peptides derived from both insulin and predicted pancreatic proteins. Lastly, *Aire*^-/-^ mouse sera were used to identify numerous auto-antigens, many of which were also observed in previous studies of humans with autoimmune polyendocrinopathy syndrome type 1 (APS1) carrying recessive mutations in AIRE. Among these were peptides derived from Perilipin-1, a validated autoimmune biomarker of generalized acquired lipodystrophy in humans. Autoreactivity to Perilipin-1 correlated with lymphocyte infiltration in adipose tissue and underscores the approach in revealing previously unknown specificities. These experiments support the use of murine proteome-wide PhIP-seq for antigenic profiling and autoantibody discovery, which may be employed to study a range of immune perturbations in mouse models of autoimmunity.

## SIGNIFICANCE

A breakdown in immune tolerance to self-antigens leads to autoimmune diseases, which are frequently characterized by the presence of autoantibodies. Defining the specificity and origins of the autoantibody responses is key for developing diagnostics and therapeutics for this family of diseases. We present a facile, inexpensive, and programmable approach for identifying global autoantibody reactivities to the murine proteome. Leveraging genetic manipulations available in the mouse model, we demonstrate the sensitivity and specificity of the PhIP-seq approach using mice lacking antibodies or those with restricted repertoires. We define and orthogonally validate a suite of known and novel autoantigens in three distinct models of autoimmune disease. Our data indicate that murine PhIP-seq can be utilized to build a more comprehensive view of autoreactive landscapes in these important models of human disease.

## INTRODUCTION

Autoimmune diseases arise from a complex breakdown in immune tolerance and are frequently characterized by the presence of autoantibodies and autoreactive T cells. Autoimmunity spans a breadth of different clinical subtypes and patterns with almost any organ system or tissue being susceptible. Defining the specificity and origins of the autoimmune response is key for developing methods to diagnose, prevent, and treat this family of diseases. On a mechanistic level, a key tool for unraveling autoimmunity has been the use of mouse models of human autoimmune diseases. A combination of autoimmune-susceptible mouse strains or mouse lines with models of human genetic defects has played a key role in our understanding in the pathogenesis of an array of autoimmune diseases. For example, use of genetically altered mice has allowed for our understanding of how the monogenic autoimmune diseases autoimmune polyendocrine syndrome type 1 (APS1)(1) and IPEX(2, 3) are linked to defects in thymic central tolerance or the function of T regulatory cells, respectively. Given the highly controlled nature of mouse modeling for both environment and genetic influences, it remains an essential tool for dissecting autoimmunity. An important aspect to this work is defining the autoimmune response in these mouse models. In this regard, a typical approach has been to search for autoantibodies from affected mice in targeted assays such as western blotting and indirect immunofluorescence. Recently, there has been rapid development of new approaches to identifying autoantibody specificities that broadly cover the entire proteome in the human setting(4–9). Thus, a similar approach in the mouse model could serve as an important method to further define and unravel autoimmunity.

Phage display immunoprecipitation and sequencing (PhIP-seq) is a powerful tool to identify antibody targets, originally described by Larman, Elledge, and colleagues(4, 10). Since 2011, it has been used to discover novel antibody autoreactivities in a wide range of human syndromes, including paraneoplastic diseases(6, 7) and inborn autoimmune syndromes(8, 9, 11). In PhIP-seq, libraries of long oligonucleotides encoding overlapping peptides are synthesized as DNA oligomers and cloned into the T7 phage genome. These libraries are expressed fused to Gene-10 of the surface-exposed capsid protein on lytic T7 phage and used as bait for antibodies in patient sera. Complex, multi-antigen immunoprecipitants are deconvoluted by sequencing the enriched phage-encoded peptides to identify multiple antibody targets in a single reaction.

Due to the programmable nature of PhIP-seq, any proteome may be comprehensively encoded into a phage library in principle. Library designs may also be highly customized, including coverage of specific protein isoforms, putative coding regions, and other features. Here, we present the construction and validation of a murine proteome-wide PhIPseq library, based on the GRCm38.p5 *Mus musculus* genome, comprised of over 480,000 peptides, representing over 76,000 protein sequences. Taking advantage of genetic manipulations available in the mouse model, library performance was evaluated across seven mouse strains, including *Rag2*^-/-^, μMT, C57BL/6J (wildtype strain “B6”), OB1, *Lyn*^-/-^, polygenic non-obese diabetic (NOD), and *Aire*^-/-^. Mice lacking mature B cells (*Rag2*^-/-^ and μMT) were used to determine proteome-wide background binding in immunoprecipitations (IPs), while serum from B6, OB1, *Lyn*^-/-^, NOD, and *Aire*^-/-^ mice was used to identify strain-specific autoreactivities. Building upon our previous identification of autoantibodies to Perilipin-1 in *Aire*^-/-^ mice(9), we identified the binding epitope of these antibodies and demonstrate their relationship to immune cell infiltrates in adipose tissue in affected mice. Taken together, these results demonstrate the utility of the approach for a broad assessment of the array of autoimmune specificities in various mouse models.

## RESULTS

### Design and construction of murine proteome-wide library

To construct a *Mus musculus* proteome-wide library, the GRCm38.p5reference proteome sequences, including all isoforms, were downloaded from NCBI and divided into 62 amino acid peptide tiles with 19 amino acid overlaps (**Fig 1A**). The library was supplemented with several positive and negative control peptides, including those derived from the human glial fibrillary acid protein (GFAP), human tubulin, GFP, and others. The resulting library of 482,672 peptides was synthesized (Agilent, Inc) as a DNA oligomer pool (**Fig 1A**) and cloned into T7 phage fused in-frame with Gene-10, the capsid protein of T7 phage. The complete peptide design file and further details are freely available as a companion to this manuscript on protocols.io (see Methods). The synthesized oligomer library and the packaged library were sequenced by next generation sequencing (NGS) with 173 and 79 million paired-end 147 base pair reads on an Illumina NovaSeq6000, respectively, which resulted in an approximate 360x and 163x coverage of the library, respectively. Alignment of the reads from sequencing of the pooled DNA oligomer library yielded 89.2% identical matches, with greater than 99.9% of all the expected peptides represented. Sequencing of the packaged T7 phage library yielded an alignment rate of 78.9%, yet representation remained high at greater than 99% of the expected peptides (**Fig 1B**).

**Figure 1.**
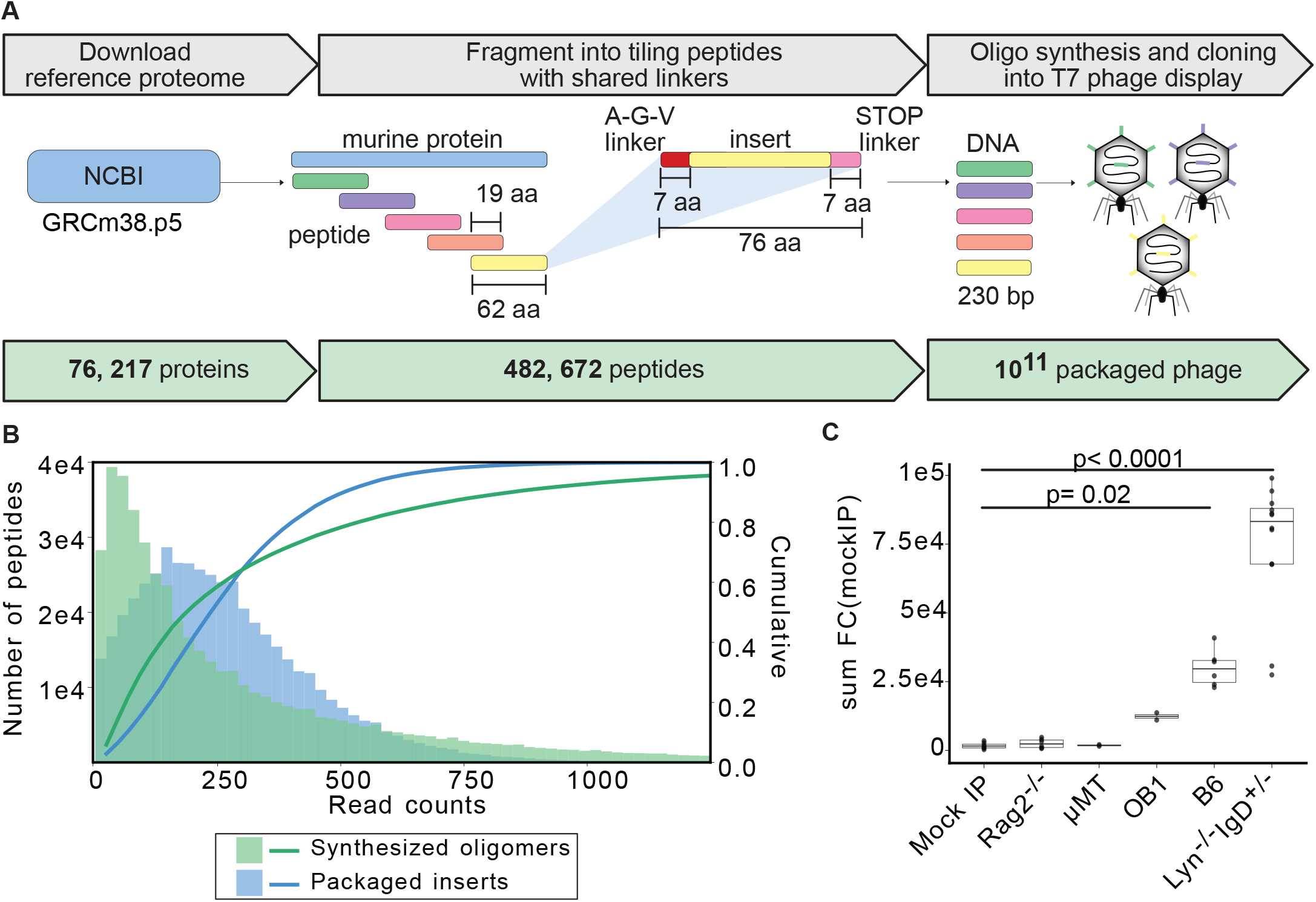
Design and validation of murine PhIP-seq library. **A.** GRCm38.p5 annotated proteins were downloaded from Refseq and 62 amino acid (aa) tiles were chosen to cover the 76,217 proteins with 482,672 peptides with a 19 aa overlap. The tiles contained necessary cloning sites for expression in T7 phage display system. **B.** Representation of designed oligos after oligo synthesis and cloning. **C**. Sum of all fold changes above the mean read counts in mock-IP in each experimental sample (mock IP) or mouse strain (*Rag2*^-/-^, μMT, OB1, B6, and *Lyn*^-/-^ *IgD*^+/-^) by PhIP-seq. Exact p-value is reported, each dot corresponds to a mouse or mock-IP replicate; Kruskal Wallis test with Tukey HSD post-hoc.

### Evaluation of library performance and background binding in Rag2^-/-^ and μMT mice

The performance of the packaged library was first benchmarked utilizing a commercial antibody with a known specificity. Previously, we have utilized commercial anti-GFAP polyclonal antibody as a positive control for human PhIP-seq libraries, due to its consistent immunoprecipitation performance (8, 11–13). The murine PhIP-seq library contains both human and mouse GFAP sequences and binding to sequences of both species was expected with the commercial antibody. Antibody bound to a mix of Protein A and Protein G magnetic beads was used to immunoprecipitate phage from the murine library, followed by either sequencing or further amplification in *Escherichia coli*. As observed previously with the human PhIP-seq library(8, 13), additional rounds of phage immunoprecipitation followed by amplification in *E. coli* resulted in increased enrichment of the target sequences. Using two rounds, approximately 20% of resulting phage encoded peptides derived from either human GFAP or mouse *Gfap* (**Fig S1A**). Using three rounds, the enrichment approached 50%, or 1x10^8^-fold greater than the amount of the same phage in the starting library (**Fig S1A**). Given the significant enrichment of *Gfap* peptides relative to non-specific peptides, all subsequent experiments utilized three rounds of immunoprecipitation and amplification.

The library was next evaluated across five mouse strains on the C57BL/6J (B6) background. Two strains (*Rag2*^-/-^ and μMT) lacking IgG(14, 15) were utilized to evaluate background binding. When compared to a mock IP control lacking serum, sera from both *Rag2*^-/-^ and μMT mice failed to significantly enrich phage from the murine library, as expected (**Fig 1C**, **Fig S1B**). At the individual peptide level, fewer than 10 peptides were consistently enriched (at least 2 of 3 mice) by sera from either mouse strain, presumably through non-specific interactions (**Fig S1C**). For subsequent experiments, the mean frequency of each phage across mock IP, *Rag2*^-/-^, and µMT was determined and used to calculate fold-change and z-scores for experimental samples (murine background model, “MBM”).

B cells in OB1 mice harbor physiological BCR rearrangements in the IgH and Ig*k*; locus that encode for a clonal IgG1 receptor with specificity for the chicken ovalbumin protein, thus their sera contains predominantly high affinity anti-ovalbumin IgG1 antibodies (16). Unlike *Rag2*^-/-^ and µMT mice, sera from OB1 mice yielded a moderate amount of enrichment consistent with the restricted B-cell repertoire of this genetic strain (**Fig 1C**, **Fig S1B**). In contrast, sera from wildtype B6 mice yielded significant enrichment with 3.2-fold more enrichment than OB1 sera (**Fig 1C**, **Fig S1B**). Finally, *Lyn*^-/-^ *IgD*^+/-^ mice were also examined, which exhibit polyclonal B-cell activation and develop lupus-like disease(17, 18). Consistent with a greater degree of autoreactivity in these mice, *Lyn*^-/-^ *IgD*^+/-^ sera yielded the largest number of significantly enriched peptides with higher fold-change than sera from wild type B6 mice (**Fig 1C**, **Fig S1B**).

### Proteome-wide PhIP-seq identifies OB1 reactivity to known epitopes in mouse proteome

Previously, the recognition site for immunoglobulin binding to the ovalbumin protein in the OB1 strain was determined as “DKLPGFGDSI” by alanine mutagenesis, where the Phe-Gly-Asp (FGD) sequence was essential for BCR binding(16). Although chicken ovalbumin sequences were not included in this murine PhIP-seq library, over 2,000 peptides in the library contain sequence similarity (p < 0.0001; see Methods) to the ovalbumin target sequence, and of those, 241 contain the critical Phe-Gly-Asp core sequence. Furthermore, an additional 2,028 peptides contain the core Phe-Gly-Asp, but lack similarity in the regions flanking this motif (see Methods, **Supplementary Table 1**).

Using sera from OB1 mice, 193 peptides were enriched greater than 4-fold above the average of MBM with a z-score of greater than three (**Fig 2A**). To investigate similarity of enriched peptides to known OB1 BCR recognition epitope, a scoring system for each peptide was developed using the sum of Smith-Waterman alignment scores to Phe-Gly-Asp across 4 amino acid sliding windows and weighted if exact Phe-Gly-Asp match was present (see Methods). Most peptides enriched by OB1 sera exhibited high epitope similarity scores, indicating specific enrichment of peptides containing the known OB1 BCR binding site (**Fig 2A**). Visual inspection of the most enriched peptides revealed significant sequence similarity to the known ovalbumin target sequence, including the core Phe-Gly-Asp (**Fig 2B**). We noted a Hnrnpa2b1 peptide containing two Phe-Gly-Asp repeats among the most highly enriched, indicating that valency within the peptide may contribute to greater enrichment.

**Figure 2.**
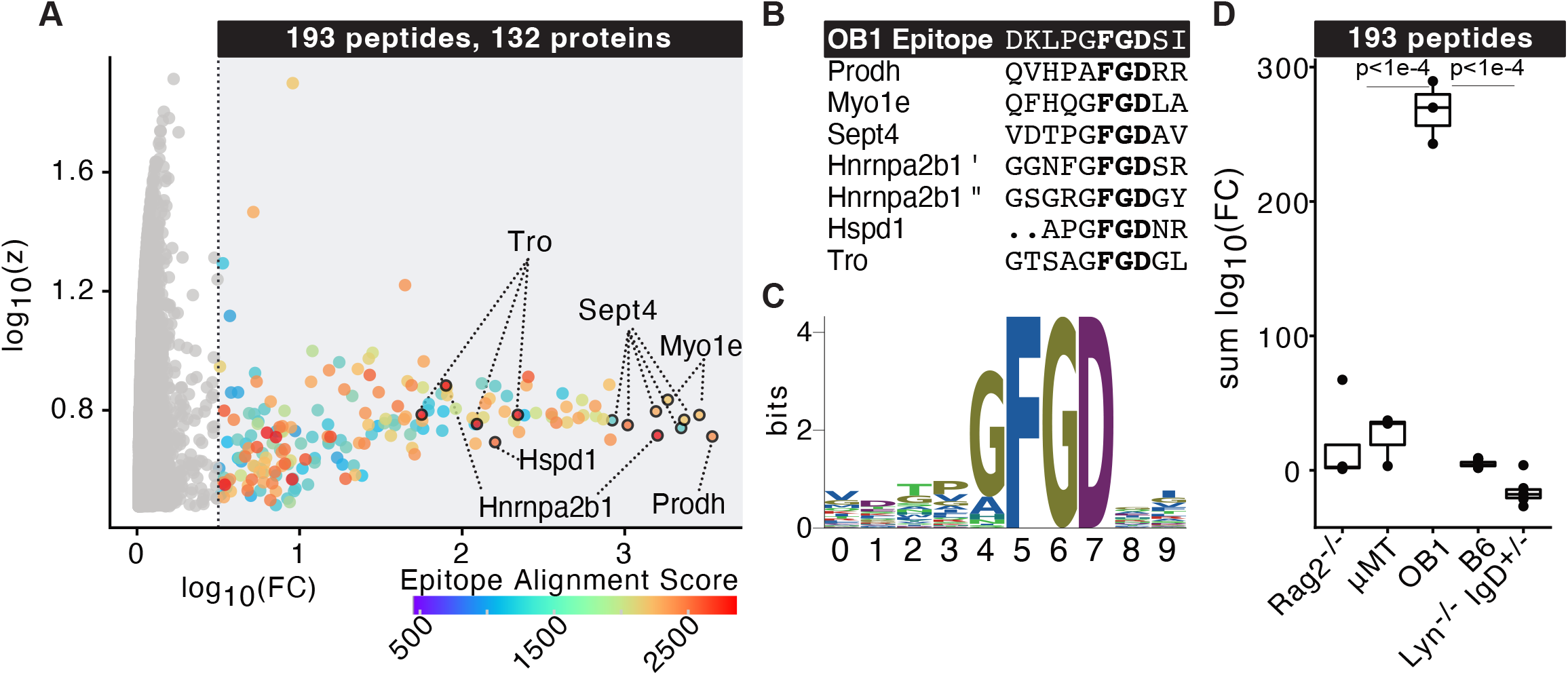
Identification of autoreactive epitopes recognized by ovalbumin-specific BCR transgenic (OB1) mice. **A.** Log_10_ fold change and z-score over murine background model (mean of *Rag2*^-/-^, μMT, and mock IP) of peptides enriched by PhIP-seq in OB1 mice colored by alignment score to known epitope and essential FGD motif. **B**. Multiple-sequence alignment of top OB1-enriched peptides **C**. Logoplot of multiple sequence alignment of 193 peptides enriched by OB1 sera. **D**. Sum log_10_ fold change over MBM of OB1 peptides enriched by sera from *Rag2*^-/-^, μMT, OB1, B6, or *IgD*^+/-^ *Lyn*^-/-^ mice. Exact adjusted p-value is reported, each dot corresponds to a mouse; Kruskal Wallis test with Tukey HSD post-hoc.

Enriched peptides containing the core sequence were aligned, revealing over representation of N-terminal Gly (**Fig 2C**), suggesting that this residue may contribute to recognition. Overall, 80% of the significantly enriched peptides contain the Phe-Gly-Asp core recognition sequences. Compared to *Rag2*^-/-^, µMT mice, wildtype, and *Lyn*^-/-^ *IgD*^+/-^ mice, these 193 significantly enriched peptides from the OB1 mice were highly specific to this strain (**Fig 2D**).

### Lyn^-/-^ IgD^+/-^ mice exhibit lupus-like autoantibody reactivity to nuclear and apoptotic antigens

Deficiency of the Src family kinase *Lyn* results in widespread autoantibody production and lupus-like disease because *Lyn* plays a non-redundant negative regulatory role downstream of the BCR by mediating ITIM-dependent inhibitory signaling; such autoimmunity is accelerated in *IgD* knockout heterozygotes (17, 19). Given the profound defect in self-tolerance in this model, we further probed sera from this model across our library. Sera from *Lyn*^-/-^ *IgD*^+/-^ mice (n=6) and wildtype B6 mice (n=6) were used to immunoprecipitate phage from the mouse PhIP-seq library. Sequencing libraries were prepared after three rounds of immunoprecipitation and amplification. The resulting read counts for each peptide were normalized to the total number of reads in each library and then averaged across all technical replicates for each mouse. High correlations (>0.75 Pearson r) were observed between technically replicated samples (**Fig S1D**). A stringent set of requirements was then used to identify peptide enrichments that were specific to the mutant mice relative to controls. A total of 508 peptides, derived from 425 proteins (**Supplementary Table 2**; **Fig S2A**), had a fold-change of at least 2-fold, a z-score of least 3, and were not enriched in any of the control B6 mice immunoprecipitations.

Approximately 37% of mutant specific peptide enrichments corresponded to proteins known to be components of or related to nuclear proteins (**Fig 3A**), consistent with the high degree of anti-nuclear staining observed in these mice (**Fig S3**) and in patients with lupus(20). To validate these findings, *Lyn*-specific PhIP-seq reactivities were compared to results from a 96-autoantigen array probed with sera from *Lyn*^-/-^ mice (**Supplementary Table 3**). Among protein autoantigens represented in the array (n=63), 32 proteins were significantly enriched in *Lyn*^-/-^ IgG above wildtype controls (see Methods), 20 of which were also identified by PhIP-seq in *Lyn*^-/-^ *IgD*^+/-^ mice (**Supplementary Table 3**). Orthogonally validated *Lyn*-specific reactivities were those to nuclear antigens such as nuclear proteins (SP100), complement (C1q), collagen (Collagen VI) and laminin (Lama1), as well as additional nuclear proteins (**Fig S2B-D; Supplementary Table 3**). In addition to sRNPs, PhIP-seq identified proteins related to their generation which were not included in the targeted autoantigen array (**Fig 3B**). This over representation of sRNPs autoimmune targets in *Lyn*^-/-^ *IgD*^+/-^ mice is consistent with prior reports that CD72-sRNP negative signaling in B-cells is *Lyn*-dependent (21, 22). Identification of nucleic-acid associated autoantigens is also consistent with an obligate role for nucleic-acid sensing machinery in B cells in this mouse model(23) and in patients(24–26). PhIP-seq also identified 14 E3 ubiquitin ligases or E3 ubiquitin complex proteins and 50 additional proteins related to apoptosis (**Fig 3C**), which is notable given evidence of impaired clearance of apoptotic debris in lupus patients(27, 28).

**Figure 3.**
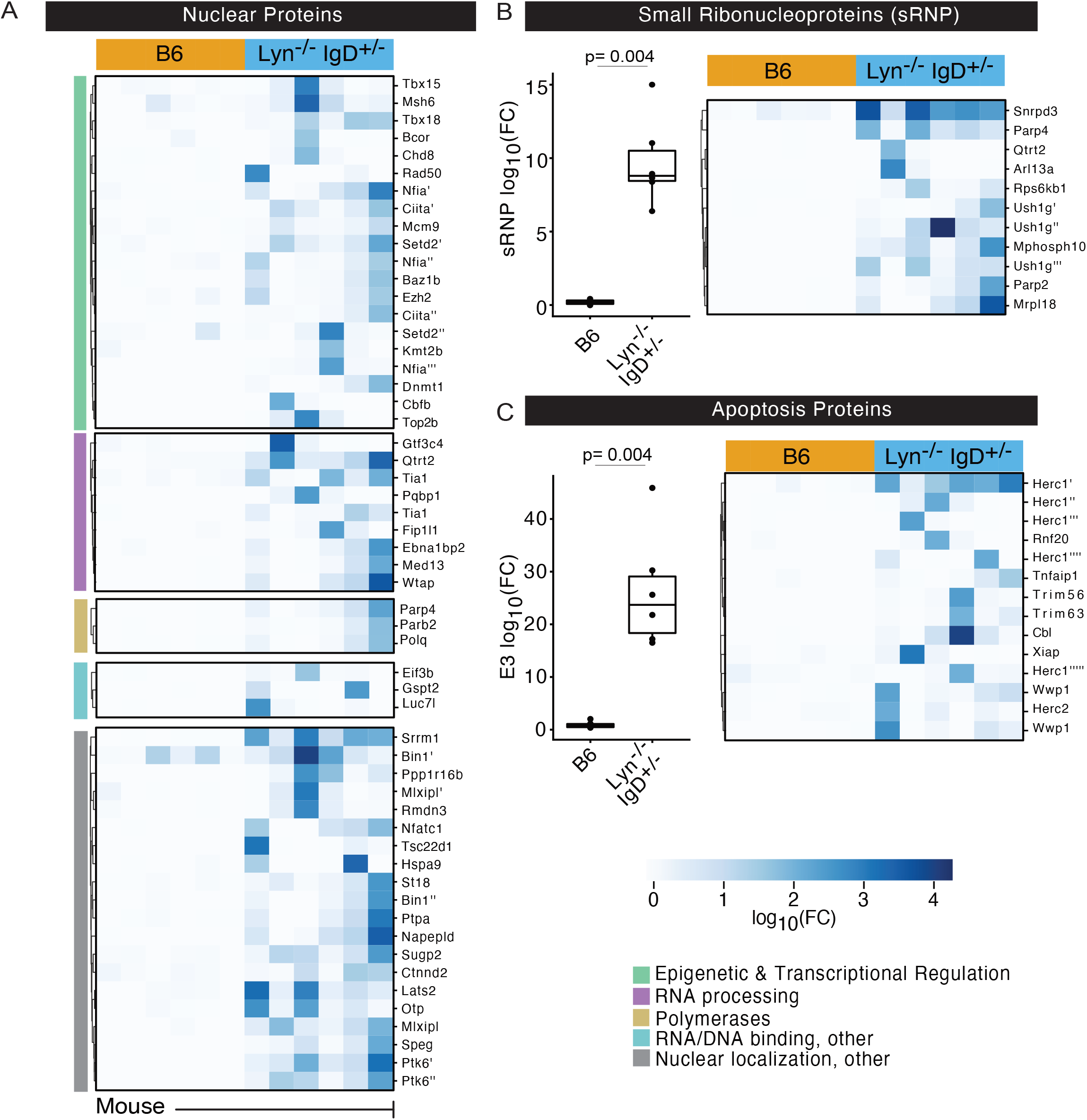
Autoreactivity to nuclear and apoptotic antigens in *Lyn*^-/-^ *IgD*^+/-^ mice. **A**. Heatmap of nuclear proteins enriched in *Lyn*^-/-^ *IgD*^+/-^ versus B6 mice. **B.** Sum log_10_ fold change over mean background (left) and heatmap of log_10_ fold change (right) of small ribonucleoproteins in B6 or *Lyn*^-/-^ *IgD*^+/-^ mice. **C**. Sum log10 fold change over mean background (left) and heatmap of log10 fold change (right) of E3 ubiquitin ligases in B6 or *Lyn*^-/-^ *IgD*^+/-^ mice. Peptide enrichments were identified by PhIP-seq in A-C. Exact adjusted p-value is reported, each dot corresponds to a mouse; Kruskal Wallis test with Dunn post-hoc.

### Autoreactivities in non-obese diabetic (NOD) versus B6 mouse strains

The murine PhIP-seq library was next used to interrogate autoantibody profiles in polygenic NOD mice (n=9), C57Bl/6 (B6) mice (n=11). NOD mice are an inbred strain that are predisposed to spontaneous autoimmunity that includes autoimmune diabetes. These mice carry an I-Ag7 MHCII haplotype that contributes to the generation of autoreactive T cells (29–32) and the development of autoantibodies with age, including those targeting insulin in older mice (33, 34). Using a z-score threshold of three relative to the MBM, modest but significant enrichment of peptides derived from mouse insulin (Ins1) and insulin-related proteins (Irs4, Insrr, Ide, Insm1) were observed (**Fig S4A-B**) in NOD mice, but not in B6 mice.

Using the stringent enrichment criteria established for the *Lyn*^-/-^ *IgD*^+/-^ experiment, a total of 314 putative autoantigens were identified (**Supplementary Table 4**). These included proteins expressed by pancreatic islet cells (Ptch1, Disp1, Pck1, Cacna1e) as well as proteins known to be associated with diabetes mellitus susceptibility or glucose metabolism (Adcy8, Perm1, Sorbs1, Pla2g6; **Fig S4B-C**). NOD mice develop additional autoimmune phenotypes such as Sjorgen’s (affecting salivary and lacrimal glands)(35), prostatitis (affecting seminal vesicle and prostate)(36), and respiratory infiltrates in older mice(37). Autoantigens affecting these tissues were also observed in NOD mice (**Fig S4C**). For example, seminal vesicle proteins Svs3 and Svs4, were enriched in male NOD mice (**Fig S4C**) consistent with reports that antibodies to seminal vesicle proteins are associated with lymphocytic infiltration of the prostate (38).

### Identification of autoreactivity and epitope mapping of anti-Plin1 antibodies in Aire^-/-^ mice

We next investigated mice lacking the transcriptional regulator *Aire*, which drives expression of tissue-specific antigens in the thymus and is required for selection of the self-tolerant T-cell repertoire (1). In humans, mutations in the AIRE transcription factor result in autoimmune polyendocrine syndrome type 1 (APS1), a rare monogenic disease in which patients develop multi-organ autoimmune pathologies due to autoreactive T cells and high-affinity tissue-specific B-cells (39). Using a human proteome-wide PhIP-seq library, our lab has previously characterized APS1 patient sera and identified a wide array of novel autoreactive antigens (8, 9, 11).

To assess proteome-wide autoreactivity in *Aire*^-/-^ mice, we performed PhIP-seq with sera of these mice on the NOD background and identified enriched peptides with a z-score greater than three above MBM. Around 57% of the known autoimmune targets in human APS1 (n=30 proteins) (11) were orthologous to putative autoreactive targets in *Aire*^-/-^ mice and around 40% of genes with known *Aire*-dependent thymic expression (40) were among *Aire*^-/-^-specific reactivities by PhIP-seq (**Fig 4A**). Using the same stringent criteria as the *Lyn*^-/-^ *IgD*^+/-^ experiment, serum from *Aire*^-/-^ but not control mice enriched 861 peptides, representing 698 proteins, which was significantly more than *Lyn*^-/-^ *IgD*^+/-^ mice (861 versus 494; **Supplementary Table 5; Fig S5A**). These included Plin1 and Muc5b (**Fig 4B-C**), which were both detected in APS1 patients and their expression is dependent on *Aire* in the thymus. Using a 3-fold cross validated logistical regression model trained on control and *Aire*^-/-^ PhIP-seq data, *Plin1* (Perilipin-1) was the top-ranked feature by the logistic regression coefficient (**Fig 4D**). Plin1 was previously identified as an autoimmune marker of generalized lipodystrophy in both mice and humans, including at least one patient with APS1(9, 41).

**Figure 4.**
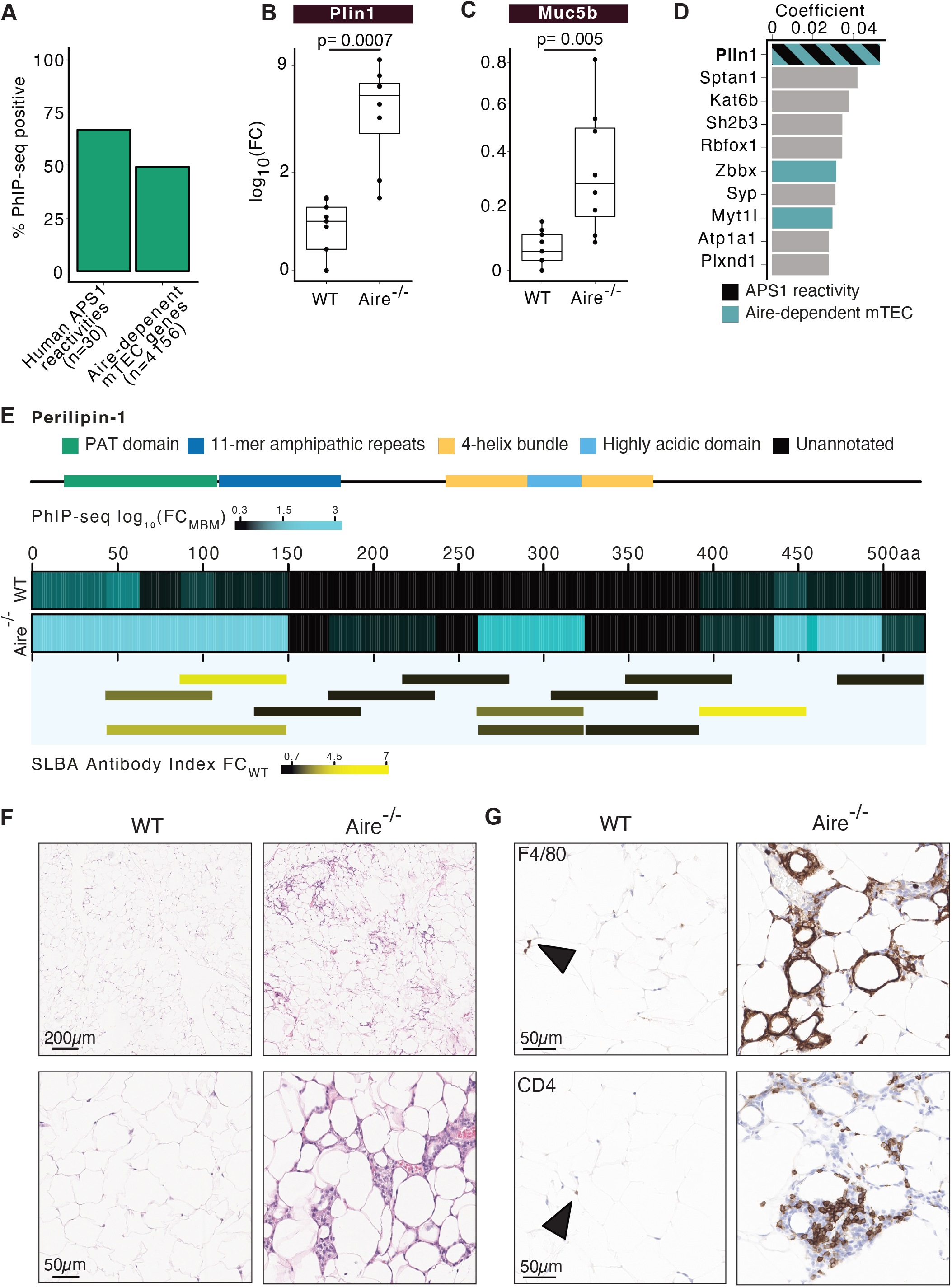
Autoreactivity in *Aire*^-/-^ mice. **A.** Percent of peptides enriched in NOD.*Aire*^-/-^ mice by PhIP-seq compared to orthologues of previously reported APS1 reactivities in humans (11) and genes under the control of *Aire* in mTEC (40). Sum log_10_ fold change over murine background model of **B.** Plin1, **C.** Muc5b in *Aire*^-/-^ versus control mice. **D.** Logistical regression coefficients of top 10 proteins for classifying *Aire*^-/-^ versus control mice colored by orthologous APS1 reactivity and/or *Aire*-dependent mTEC expression. **E**. Heatmap of perilipin 1 positional sum log_10_ fold change over background in *Aire*^-/-^ or wildtype mice by PhIP-seq annotated with domain positions (top). Fold change of antibody index in *Aire*^-/-^ over wildtype mice by SLBA (bottom). **F.** Inguinal fat pads stained with H&E, higher magnification below. **G.** Immunohistochemistry of F4/80 (top) or CD4 (bottom) in inguinal fat pads in *Aire*^-/-^ versus NOD mice. Arrowhead indicates positive cells. Peptide enrichments were identified by PhIP-seq in A-E; SLBA in E (bottom). Exact adjusted p-value is reported, each dot corresponds to a mouse; Kruskal Wallis test with Dunn post-hoc.

Examination of the peptide level enrichment data for Plin1, across all eight *Aire*^-/-^ mice revealed differential enrichment in the PAT domain, and to a lesser extent, the 11-mer amphipathic repeat region, and the highly acidic domain (**Fig 4E**). The PAT domain is highly conserved in the perilipin family of proteins, yet other perilipins (*Plin2*, *Plin3*, *Plin4*, and *Plin5*) did not exhibit enrichment in *Aire^-/-^* mice (**Fig S5B**), suggesting that the epitope is specific to *Plin1*. Positional conservation analysis of the PAT domain across perilipins showed greatest divergence in two regions corresponding with highest *Aire*^-/-^ reactivity within Plin1 (**Fig S5C**), plausibly explaining antibody specificity to Plin1 but not other members of the family.

To validate these findings, a split luciferase binding assay (SLBA) was developed by generating peptides (n=13) tiling across Plin1 with a HiBiT tag that complexes with LgBiT to produce luminescence (**Fig 4E**; **Fig S6A**). SLBA possess several advantages for rapid orthogonal validation of PhIPseq results, including features of the luciferase immunoprecipitation system (LIPS)(42) and the radioligand binding assay (RLBA)(43). All Plin1 fragments generated at least five-logs greater luciferase signal than an in-frame stop codon construct when LgBiT and luciferase substrate was added (**Fig S6B**). IPs were performed with tagged peptides and *Aire*^-/-^ or control sera. Luciferase activity was measured after IP and fold-change over wildtype control mice was calculated for antibody indices using an anti-HiBiT antibody (see Methods). The most highly enriched SLBA fragments corresponded to the most enriched PhIP-seq peptides in the PAT domain, with the exception of one enriched fragment in a C-terminal unannotated region (**Fig 4E**).

As noted, anti-PLIN1 is a known a marker for acquired generalized lipodystrophy in humans(9). Generalized lipodystrophy is characterized by progressive loss of adipose tissue, panniculitis, and metabolic abnormalities(44). To investigate whether *Aire*^-/-^ mice also exhibit similar features, gonadal fat pads were analyzed by H&E staining (**Fig 4F**). *Aire*^-/-^ mice exhibited elevated cellular infiltration as compared to wildtype controls (**Fig 4F**, **Fig S7**) and immunohistochemistry analysis indicated the presence of macrophages (F4/80^+^ cells) and CD4^+^ T cells (**Fig 4G**). These data suggest that *Aire*^-/-^ driven autoimmune lipodystrophy in mice possess similar features to their human disease counterpart, with Plin1 autoreactivity being the hallmark in both.

## DISCUSSION

The PhIP-seq technique has been used extensively for the identification of autoreactivies in human sera and cerebrospinal fluid, including for the identification of novel autoimmune diseases (4–11, 13). While the technique has proven useful in human clinical investigations, a parallel murine PhIP-seq library provides new avenues to investigate immune dysregulation and disease in a tractable model organism. Here, we describe the construction and validation of a proteome wide murine PhIP-seq library, composed of greater than 480,000 peptides, each 62 amino acids in length, spanning over 76,000 protein sequences. While similar in design, this library includes approximately 27,000 additional protein sequences (including variants) than a previously published library(45). For any newly designed PhIP-seq library, it is essential to quantify the performance characteristics. Here, this murine PhIP-seq library was extensively validated using seven different mouse strains, including three well-characterized autoimmune models, as well as wildtype, OB1, and mature B cell-deficient mice (*Rag2*^-/-^ and µMT). The murine PhIP-seq library exhibited robust performance with control antibody and murine sera, lending further support for PhIP-seq as a reliable and sensitive method for large-scale detection of antibody-antigen interactions.

Given the large number of input analytes present in PhIP-seq experiments, there is the potential for non-specific interactions that could result in false positives. This includes non-immunoglobulin-dependent interactions, such as aggregation and protein binding to bead matrices. This problem is further compounded by naturally occurring autoreactivities, arising from the unique repertoire of antibodies present in every individual, most of which are unlikely to be associated with disease pathology(46). Recent work has demonstrated that PhIP-seq applications in humans require appropriately large numbers of healthy control samples to eliminate false-positives and autoreactivities not associated with disease(11). Model organisms afford unique opportunities to evaluate PhIP-seq performance. For example, using mice that do not produce immunoglobulins, less than ten “false-positive” autoreactivities were detected compared mock-IP controls, despite three rounds of iterative phage enrichment. This indicates that interactions detected by PhIP-seq primarily arise from immunoglobulin-antigen interactions and not spurious protein-phage binding.

The murine PhIP-seq library was designed with long 62 amino acid overlapping segments corresponding to all the known coding mouse sequences. However, it is well understood that linear epitope binding by antibodies typically involves recognition of relatively short sequences, ranging from 4 to 12 amino acids(47). It is also appreciated that antibodies may vary widely with respect to specificity, including binding of biochemically similar motifs, but non-identical sequences. This represents both a challenge and an opportunity. For any given collection of PhIP-seq enriched peptides, the challenge is deducing which motifs within the long phage-displayed peptides are interacting with immunoglobulins, and which peptides that derive from separate coding sequences are in fact being enriched by a shared motif. We have previously shown that motif finding algorithms, such as MEME(48), may be used to address this challenge by approximating the shared motifs amongst enriched peptides(13). However, the natural variation of sequences within a proteome is also an opportunity given that many subtle variations of biochemically shared motifs are likely to be inherently embedded in the library. This facet of PhIP-seq was demonstrated using OB1 mice, which possess a predominant high-affinity IgG specificity to chicken ovalbumin (OVA)(49). OVA is not a murine protein, and thus the precise OVA protein sequence is not represented in the murine library. Despite this fact, serum from these mice enriched over 193 peptides derived from over 133 proteins, in many cases at levels greater than 1000x over the background model. Motif analysis of enriched peptides revealed the canonical Phy-Gly-Asp motif for the known OVA epitope, and further highlighted the importance of a preceding Gly residue, that was not previously appreciated. In this manner, epitope-level biochemical motif definitions may arise from PhIP-seq analysis, even in libraries lacking exact matches, in addition to capturing the range of antibody cross-reactivity. This also highlights the complex nature of autoimmune interactions, wherein an antibody thought to be highly specific has the potential to interact with a multitude of similar sequences. While many of these off-target sequences may not be physiologically relevant, we note that off-target interactions have, at the minimum, the potential to complicate interpretations of immune dysregulation.

PhIP-seq has the inherent capacity to yield large numbers of putative autoreactive antigens, however orthogonal validation by separate assays can be a time-consuming, labor-intensive process. Here, we introduce the split luciferase binding assay (SLBA), which is a variant of the radio ligand binding assay (RLBA)(43) and the luciferase immunoprecipitation system (LIPS)(42). With SLBA, oligomers from the PhIP-seq library or full-length proteins can be rapidly fused to a split luciferase tag, which is a compact 11-amino acid sequence, without requiring cloning. The time from synthetic oligomer receipt to immunoprecipitation result can be a fast as two days and requires no radioactive reagents. Similar to RLBA and LIPS, this approach may be extended to investigate any protein-protein interaction by co-precipitation.

To further characterize the murine PhIPseq library, three autoimmune-prone mouse lines were used, specifically *Lyn*^-/-^, NOD, and *Aire*^-/-^. In all three models, unique patterns of autoreactivity were identified that corresponded with underlying defects in immune tolerance and reflect what is seen in humans. For example, anti-nuclear and apoptotic reactivities identified in *Lyn*^-/-^ mice are consistent with autoantigen array data, previous reports (21–23), and descriptions in lupus patients(24–28).

Likewise, in the NOD background murine PhIPseq analysis revealed enhanced autoreactivity relative to the wildtype B6 background, spanning proteins that derive from multiple tissues, including pancreas, lung, salivary, and male reproductive tissues. Of note, we did not detect strong enrichment of insulin-reactive autoantibodies in our approach which is a known autoantibody in the NOD mouse strain(50). This could be related to the improper display of tertiary or secondary structures that are critical for this reactivity and is a known limitation of our approach.

*Aire*^-/-^ mice afford an opportunity to investigate an extreme phenotype of autoimmune dysregulation due to a loss of central tolerance. *Aire* controls medullary thymic epithelial cell (mTEC) expression of self-antigens necessary for negative selection of autoreactive T cells (1). Nearly half of known *Aire*-dependent proteins expressed in mTECs were identified as autoreactivities in *Aire*^-/-^ sera by PhIP-seq. This suggests that failure to negatively select T cells to these proteins results in cognate autoreactivities. These results exemplify the direct relationship between *Aire*-regulated mTEC antigen expression and resultant antibody autoreactivity. Furthermore, the autoreactivities identified in *Aire* mice were highly orthologous to those in APS1 patients, which lack functional *Aire*: the majority of identified autoreactivities in humans were also present in NOD.*Aire*^-/-^ mice. These results point to the importance of unbiased proteome-wide autoreactivity profiling in building our understanding of the fundamental relationship of antigen presentation in the thymus and the development of central tolerance in mouse and humans. Looking forward, it will be interesting to see if other models with defects in central tolerance have an overlap in the array of autoantigens defined here.

The most significantly enriched autoreactive target in *Aire*^-/-^ mice was Perilipin-1, whose mTEC expression is also dependent on *Aire* (9, 40). Anti-Plin1 antibodies were recently identified in patients with acquired generalized lipodystrophy, APS1, and cancer immunotherapy treatment (9). We previously demonstrated that archived *Aire*^-/-^ mice on the B6 background exhibited anti-Plin1 autoantibodies and in this study, we identified that Plin1 is the most prevalent autoreactivity in NOD.*Aire*^-/-^ mice. Plin1 is an intracellular protein essential for lipid storage and lipolysis in adipocytes(51), therefore it is unlikely that the antibody acts on Plin1 within the cell. It is more plausible that the antibodies to Plin1 are not necessarily directly pathogenic, and the disease is T cell mediated, as has been demonstrated in a number of other *Aire*^-/-^ related autoantigens(52, 53). This scenario would be analogous to paraneoplastic autoimmune disease, like Anti-Hu encephalitis, wherein the effectors of inflammation are cytotoxic T cells, reactive to peptides derived from the same proteins to which the B-cell response is directed (e.g. anti-HuD)(54). Consistent with this notion, T cell infiltrates were observed in fat pads of *Aire*^-/-^ mice. Importantly, these results suggest that the NOD.*Aire*^-/-^ mouse model includes the spontaneous development of acquired autoimmune lipodystrophy. To date, no such animal model has been described for this disease and could open new avenues to further understand this important clinical autoimmune disease.

PhIP-seq is a modular and low-cost approach to identify antibody reactivities which may be broadly applied to further define autoimmunity in mouse models beyond the present study and is a powerful complement to traditional antigen screening techniques, such as IP-mass spectrometry. The mouse, as a model organism, affords unique opportunities to identify autoreactivities in a myriad of well-defined immunological perturbations such as immunization, infection, grafts, pregnancies, or tumors. Our newly defined and validated murine PhIP-seq approach can now be a powerful adjunctive approach to develop a more comprehensive and holistic view of autoreactive landscapes in these important models.

## MATERIALS AND METHODS

### Mouse Strains

Mouse experiments were performed in accordance with University of California San Francisco (UCSF) guidelines under approved protocols by the Institutional Animal Care and Use Committee. Mice were maintained in a specific pathogenic free facility at 25 degrees Celsius, ambient humidity, and a 12-hour day/night cycle.

Wildtype C57BL/6J, NOD, and *Rag2^-/-^* mice (B6.Cg-*Rag2^tm1.1Cgn^*/J)(55) mice were purchased from the Jackson Laboratory. Unless otherwise indicated, mutant strains were also on the C57BL/6J background. OB1 transnuclear mice(49) were a generous gift from H. Ploegh at Boston Children’s Hospital. *Lyn*^-/-^ *IgD*^+/-^ mice have been previously described (17) and serum containing known high titers of autoantibodies were used for experiments. NOD mice were obtained from Jackson Laboratory and NOD.*Aire*^-/-^ mice have been previously described(56). NOD control and *Aire*^-/-^ mice were greater than 12 weeks of age.

Serum was obtained via orbital bleed and allowed to clot for 45 minutes at room temperature, then centrifuged at 1500*g* for 10 min to collect the serum supernatant which was promptly aliquoted and frozen for temporary storage prior to experimentation.

### Library construction, cloning, and sequence validation

*Mus musculus* reference proteome sequences, including all isoforms, (GRCm38.p5) were downloaded from NCBI and fragmented into 62 amino acid peptide tiles with 19 amino acid overlap. Peptides were clustered to 95% similarity and low-complexity sequences were removed with the Lempel-Ziv-Welch algorithm. The resulting library of 482, 672 peptides was synonymously scrambled to prevent self-annealing at overlapping regions, codon-optimized for *Escherichia coli* and synthesized as an oligomer pool (Agilent Inc). Each peptide contained an alanine/glycine N-terminus linker (AAVVGGV) and a double-stop C-terminus linker (**AYAMA). The resultant oligomer library was cloned into a T7 phage display system (Millipore) by using primers to attach restriction enzyme sites, ligated into the T7 genome, and packaged into phage, as previously described (13).

Linker-specific de-phasing primers were used to amplify genomic insert corresponding to displayed peptide from the synthesized oligo pool and library after cloning. Amplicon libraries were indexed for next generation sequencing using the NovaSeq6000 platform (Illumina), as previously described(13). Step-by-step library construction protocol is available at dx.doi.org/10.17504/protocols.io.kxygx9144g8j/v1

### PhIPseq with murine serum

PhIP-seq was performed as previously described(8, 13) by incubating 0.5μL of mouse sera and 0.5 μL 2x antibody storage buffer (40% glycerol, 40mM HEPES pH 7.3, 0.04% (w/v) NaN_3_, 2x PBS) with 1mL of murine PhIP-seq library at approximately 1x10^11^ plaque forming units (pfu)/mL with over-head rotation overnight at 4°C. Bound immune complexes were captured with an equal mix of protein A and protein G Dynabead magnetic beads (Thermo Fisher Scientific), which recognize all mouse IgG subclasses, washed to remove non-specific binding, and inoculated into *E. coli* cultures for lytic phage amplification. Three successive rounds of amplification were performed for each experiment and the last two rounds were sequenced for each experiment. Each sera was tested in at least technical duplicate. Mock immunoprecipitations were performed after incubating the murine PhIP-seq library with 1 µL of antibody storage buffer in lieu of serum, but otherwise followed the same IP and amplification protocol. Positive control PhIP-seq designed to capture mouse and human GFAP began by incubating 0.1µg/mL polyclonal anti-GFAP antibody (Dako) with the murine PhIP-seq library.

### Anti-nuclear antibody staining

Anti-nuclear immunofluorescence was performed as previously described(23). Briefly, Kallestad HEp-2 slides (BIO-RAD) were stained overnight with 1µL of serum from either *Lyn*^-/-^*IgD*^+/-^ or wildtype mice. After washing, antibody binding was detected using an anti-mouse secondary antibody conjugated to Alexa488 dye (Thermo) and DAPI counterstain (Thermo). Cells were imaged with the Crest LFOV Spinning Disk/ C2 Confocal at 1000x magnification under identical camera exposure and laser settings for knockout and control mice. Micrograph exposure was normalized to secondary only negative control in FIJI and applied to all images at once.

### Identification of Lyn^-/-^ reactivities using an autoantigen array

Serum was shipped to UT Southwestern microarray core facility for analysis as previously described(57). Autoantigen array results were filtered for protein antigens, excluding non-protein antigens, complex antigens, or post-translationally modified antigens. *Lyn*-specific antigens were identified by utilizing the average signal from IgG normalized data. Means and standard deviation from wildtype mice were used to calculate z-scores in *Lyn*^-/-^ mice and an antigen was considered to be a hit if it met a z-score threshold of 3 and *Lyn*-specific if greater than two *Lyn*^-/-^ mice had the peptide as a hit while none of the wildtype mice did. *Lyn*-specific autoreactivities identified by autoantigen array were compared to autoreactivities identified by PhIP-seq in *Lyn*^-/-^*IgD*^+/-^ mice by comparing lists of hit peptides.

### Split Luciferase Binding Assay (SLBA)

A detailed SLBA protocol is available on protocols.io at dx.doi.org/10.17504/protocols.io.4r3l27b9pg1y/v1. Briefly, peptides tiling across Plin1 were inserted into a split luciferase construct containing a terminal HiBiT tag and synthesized (Twist Biosciences) as DNA oligomers. Constructs were amplified by PCR using 5’-AAGCAGAGCTCGTTTAGTGAACCGTCAGA-3’ and 5’-GGCCGGCCGTTTAAACGCTGATCTT-3’ primer pair. Unpurified PCR product was used as input to rabbit reticulocyte transcription translation system (Promega) and Nano-Glo HiBit Lytic Detection System (Promega Cat No. N3040) was used to measure relative luciferase units (RLU) of translated peptides in a luminometer. Peptides were normalized to 2e7 RLU input, incubated overnight with mouse sera, and immunoprecipitated with protein A and protein G sepharose beads (Millipore Sigma). After thoroughly washing beads with SLBA buffer (0.15M NaCl, 0.02M Tris-HCl pH7.4, 1% w/v sodium azide, 1% w/v bovine serum albumin, and 0.15% v/v Tween-20), luminescence remaining on beads was measured using Nano-Glo HiBit Lytic Detection System (Promega Cat No. N3040) in a luminometer. Anti-HiBiT antibody (Promega) was used as a positive control for each peptide. Antibody index was calculated by dividing RLU for each experimental condition by RLU obtained by anti-HiBiT antibody. Fold-change in *Aire*^-/-^ mice was calculated by dividing antibody index by the mean antibody index in wildtype control mice.

### Adipose tissue microscopy and quantification

Perigonadal fat pads were dissected, fixed in 10% neutral buffered formalin for 24 hrs, and sent for processing to Histowiz for hematoxylin and eosin staining as well as immunohistochemistry. Images were quantified in QuPath software(58) using the “cell detection” tool within a rectangular region of interest. The number of detected cells was normalized per 1e7 µm^2^ area.

### Bioinformatic and Statistical Analysis

Next generation sequencing reads from synthesized or cloned oligomer libraries were aligned to the designed library with bowtie2 (59), as previously described (8). To analyze peptide enrichment after PhIP-seq, reads were aligned at the protein level using RAPsearch(60), as previously described(8).

Aligned reads were normalized to 100,000 reads per k-mer (RPK) to account for varying read-depth and further normalized to the mean of *Rag2*^-/-^, μMT and mock IP controls within each experimental batch. Post-normalization batch effect was not observed. Auto-correlation between technical replicated samples was evaluated by Pearson correlation. RPK was averaged across technical replicates and strain-specific candidate antigens were identified using an implementation of PhagePy python package (https://github.com/h-s-miller/phagepy). Briefly, a z-score greater than or equal to 3 and fold change greater than 2 over background were required in test strain versus control. For experiments evaluating library performance relative to mock IP, per-animal z-scores were calculated for each peptide and filtered as above.

To assess the off-target binding of OB1 sera to peptides in the murine PhIP-seq library, every peptide in the library was scanned for sequence similarity to the OVA epitope DKLPG**FGD**SI using Fimo(61) with a p-value threshold of p< 0.0001. Peptides in murine proteome with significant sequence similarity to the OVA epitope were filtered by presence of the essential FGD sequence. Epitope scoring was developed to evaluate peptide enrichment of FGD motif as follows. Each peptide was scanned in a 4-mer sliding window and scored in each 4-mer window. The score was defined as 500 if the 4-mer contained the essential epitope ‘FGD’ or else given by the Smith-Waterman alignment score between ‘FGD’ and the 4-mer. The Smith-Waterman algorithm (implemented with scikit-bio) used a substitution matrix in which amino acid matches were scored 25, amino acid substitutions within Dayhoff category were scored 10 and amino acid substitutions outside of Dayhoff category were scored -25 and used a gap open penalty of 1 and gap extension penalty of 4. The total alignment score was taken to be the summation of scores across all 4-mer windows.

Human orthologs of NOD-specific peptides were surveyed for tissue expression in the salivary gland, seminal vesicle, prostate, pancreas, and lung using tissue expression data from the human protein atlas (https://www.proteinatlas.org/about/download). Peptides were labeled as ‘expressed’ in the given tissue type if their protein ortholog had ‘high’ level of expression as determined by immunohistochemistry of tissue micro arrays.

Kruskal Wallis test with Dunn’s multiple comparisons was used to evaluate significance between fold changes for selected proteins or pathways across mouse strains, unless otherwise indicated in the figure legend.

### Sequence Availability

Design files for murine PhIP-seq library, as well as raw and aligned PhIP-Seq data are available for download at Dryad.

## ACKNOWLEDGEMENTS

We thank members of the DeRisi for helpful discussions and JM Rackaitis for support during these studies. We thank the UCSF CALM-NIC microscopy core for use of the CREST LFOV Spinning Disk/C2 Confocal funded by the UCSF Program for Breakthrough Biomedical Research, the Sandler Foundation, Strategic Advisory Committee, and the EVCP Office Research Resource Program Institutional Matching Instrumentation Award. E.R. holds a 2022 Next Gen Pregnancy Research Grant from the Burroughs Wellcome Fund and the Eunice Kennedy Shriver National Institute of Child Health and Human Development T32 award (5T32HD098057). C.M.B. is funded by The Emiko Terasaki Foundation (Project 7027742/Fund B73335) and by the National Institute of Neurological Disorders and Stroke of the NIH (award 1K99NS117800-01). GR is funded by National Institute of Allergy and Infectious Diseases (award K08AI137209). J.L.D. is funded by a grant from Chan Zuckerberg Biohub. Contents herein are the sole responsibility of the authors and do not necessarily represent the official views of the NIH or other funding agencies.

No potential conflicts of interest relevant to this article were reported.

## SUPPLEMENTAL FIGURE LEGENDS

**Figure S1.**
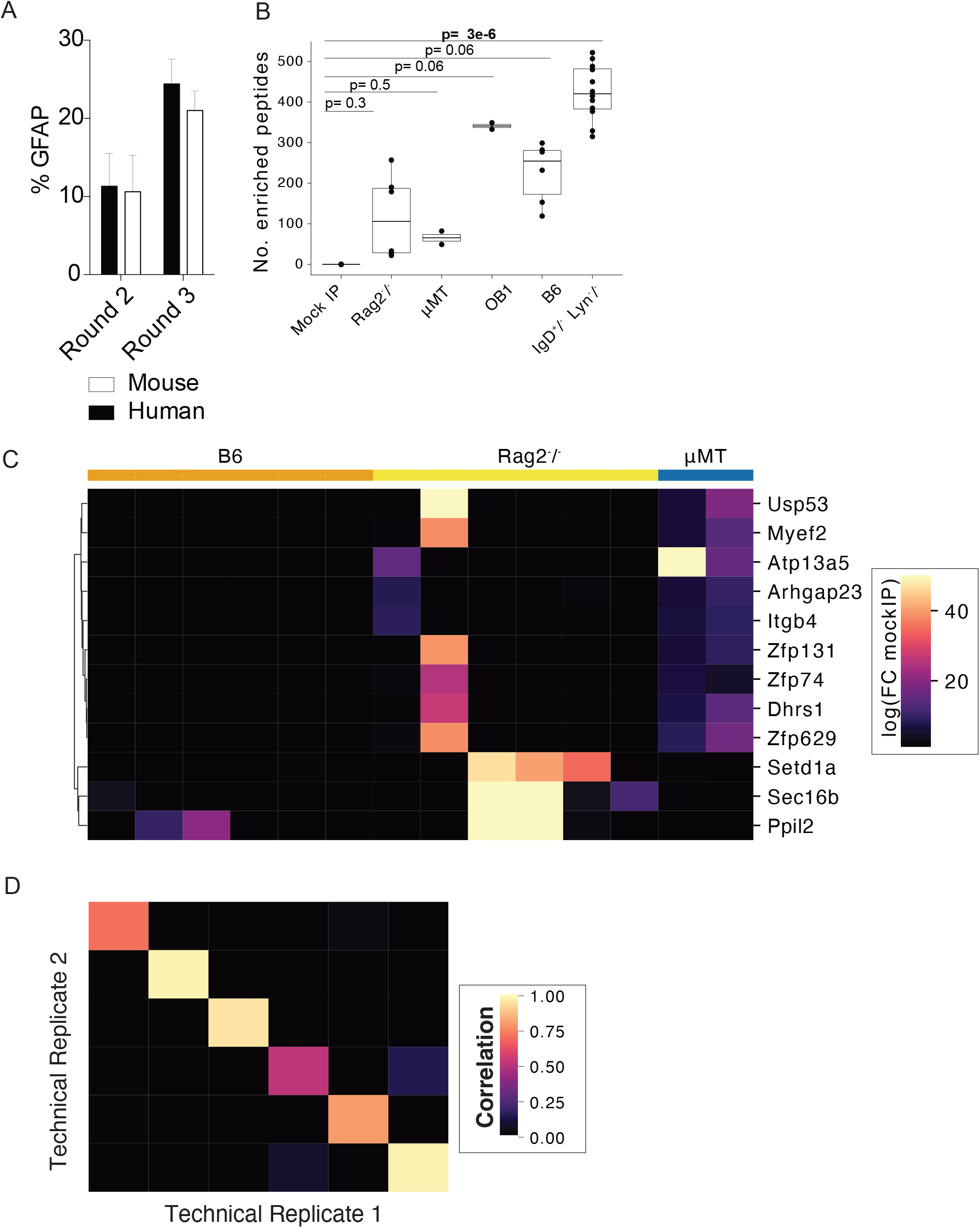
Background modeling with *Rag2*^-/-^ and μMT mice. **A.** Percent of reads mapping to human and mouse GFAP positive control peptides after two and three rounds of panning. **B.** Number of enriched peptides identified with a z-score greater than 3 and fold-change greater than 2 compared to mockIP in each experimental sample (mock IP) or mouse strain (*Rag2*^-/-^, μMT, OB1, B6, and *IgD*^+/-^ *Lyn*^-/-^). **C.** Heatmap of top peptide log 10 fold change over mock IP in *Rag2*^-/-^ or μMT mice compared to B6 mice. Exact p-value is reported, each dot corresponds to a mouse or mockIP replicate; Kruskal Wallis test with Dunn post-hoc. **D.** Heatmap of Pearson correlation of two technically replicated peptide enrichments in *Lyn*^-/-^ *IgD*^+/-^ mice by PhIP-seq.

**Figure S2.**
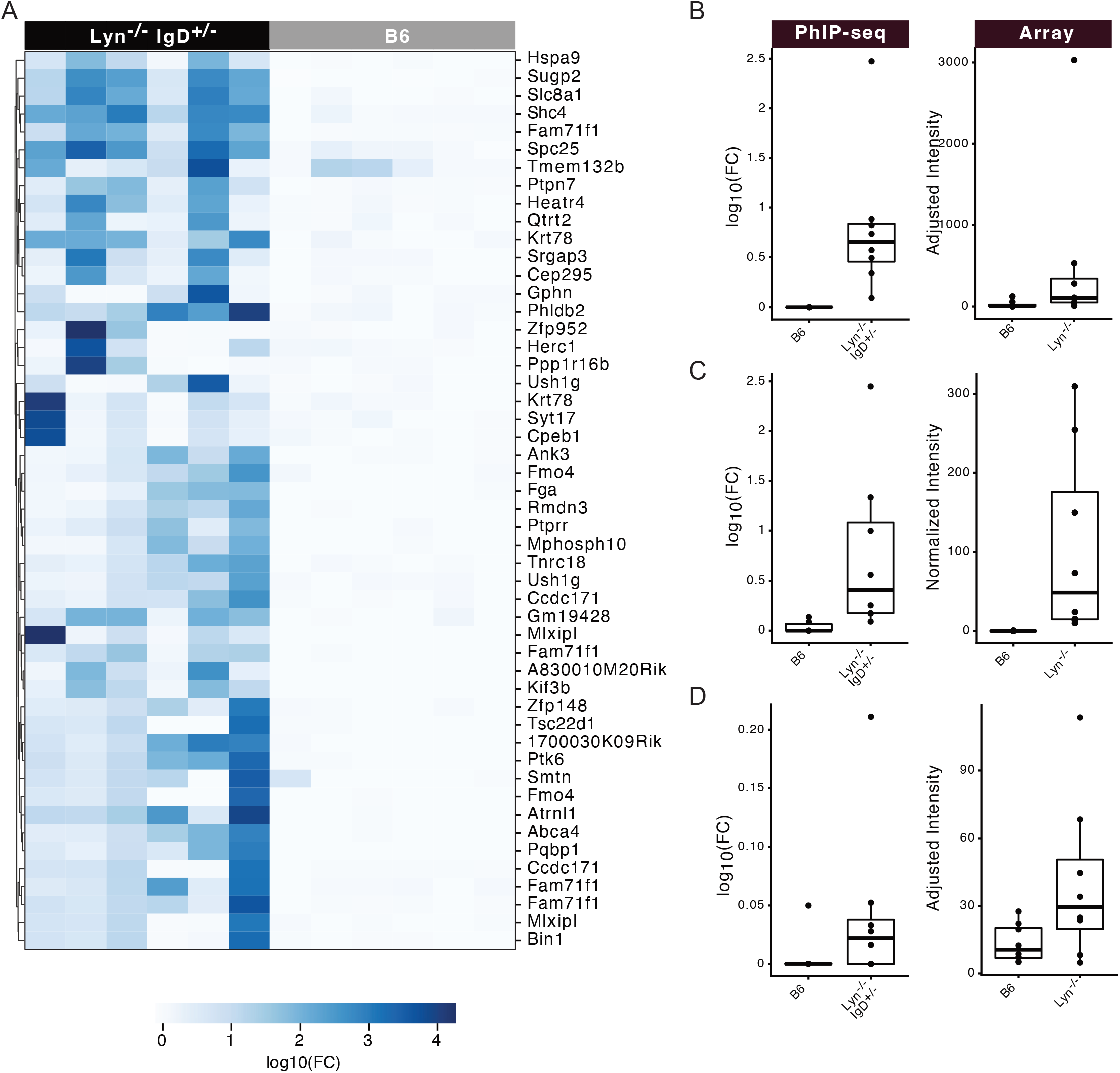
Autoreactive peptides in *Lyn*^-/-^ *IgD*^+/-^ mice. Heatmap of log_10_ fold change over MBM in **A.** top 50 peptides ranked by fold change in *Lyn*^-/-^ *IgD*^+/-^ versus B6 mice. Peptide enrichments were identified by PhIP-seq in A and annotated by their corresponding protein. Log_10_ fold change over MBM or normalized intensity of **B.** Snrp/SmD **C.** Collagen VI, **D.** or Laminin in *Lyn*^-/-^*IgD*^+/-^ (left) or *Lyn*^-/-^ (right) or wildtype control mice as detected by PhIP-seq (left) or autoantigen array (right).

**Figure S3.**
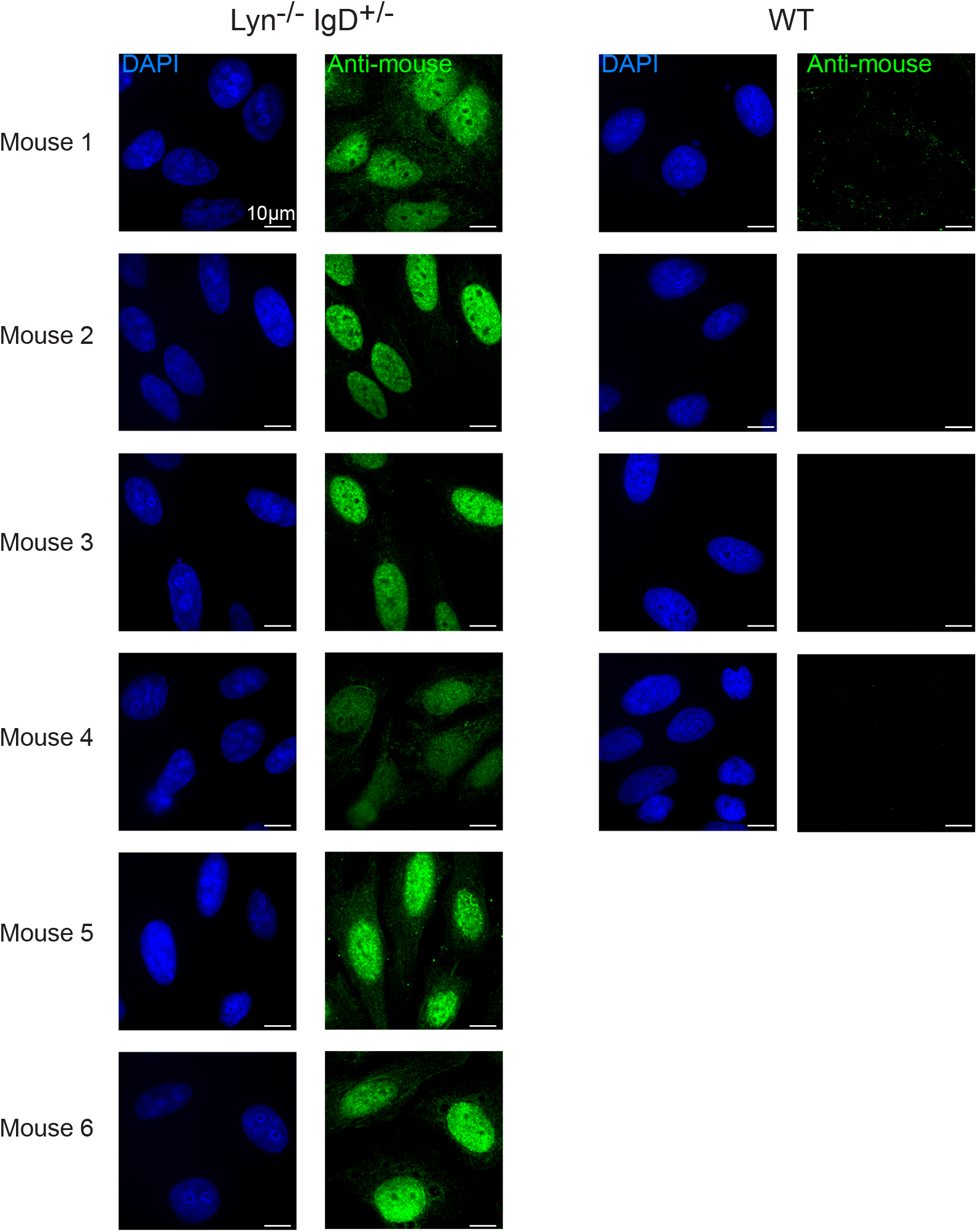
Anti-nuclear antibodies in *Lyn*^-/-^ *IgD*^+/-^ mice. Representative images of immunofluorescence detection of *Lyn*-^/-^*IgD*^+/-^ (left) or wildtype (right) mouse sera binding to nuclei of HEp-2 cell line. Side-by-side single channel emission for DAPI and anti-mouse Alexa488 is shown for each mouse. Scale bar corresponds to 10µm.

**Figure S4.**
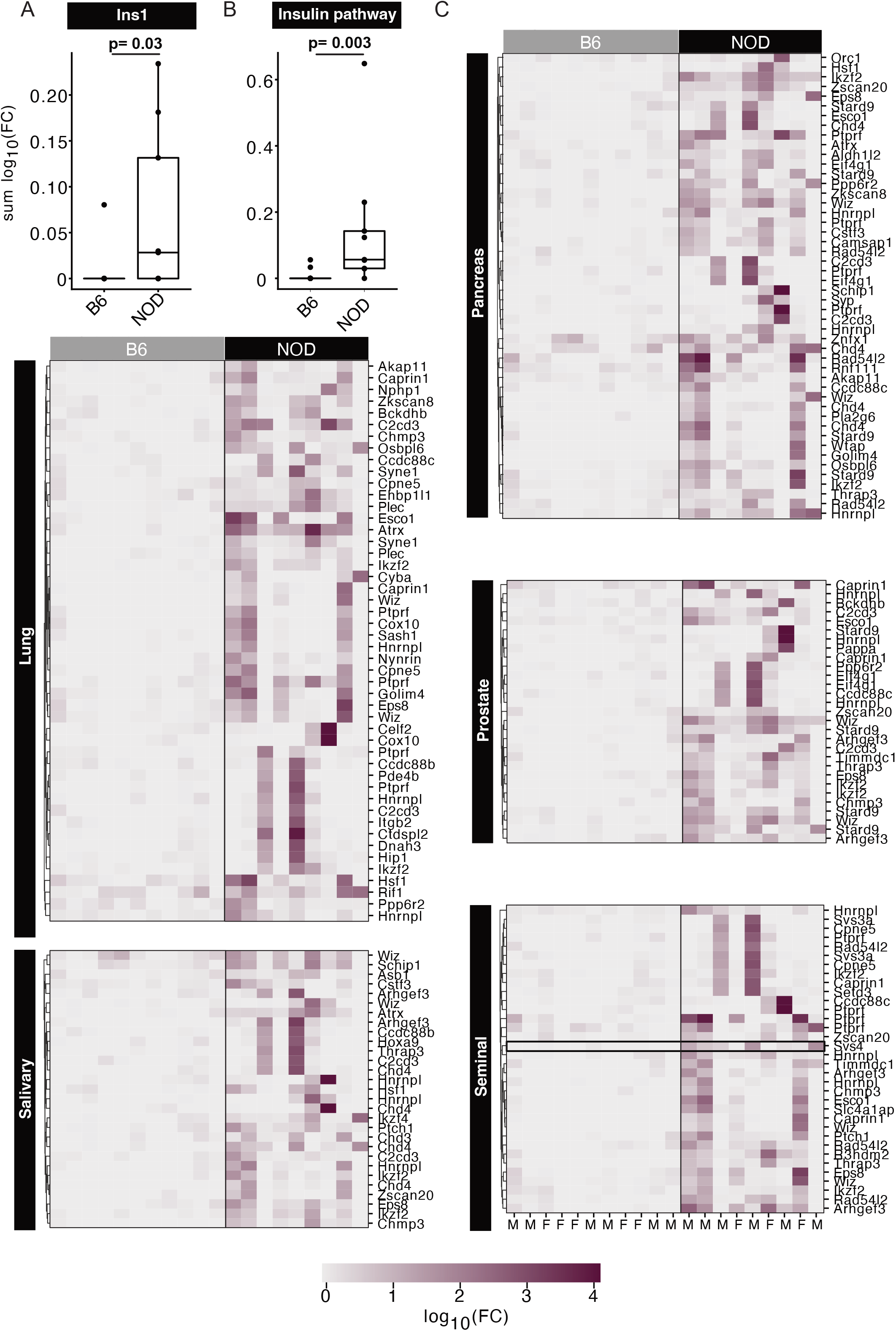
Autoreactive peptides in NOD mice. **A.** Sum log_10_ fold change over MBM and **B-C**. heatmap of log_10_ fold change of insulin pathway (A) or predicted pancreatic proteins (B) or 50 peptides ranked by fold change (C) in B6 or NOD mice. Peptide enrichments were identified by PhIP-seq in and annotated by their corresponding protein.

**Figure S5.**
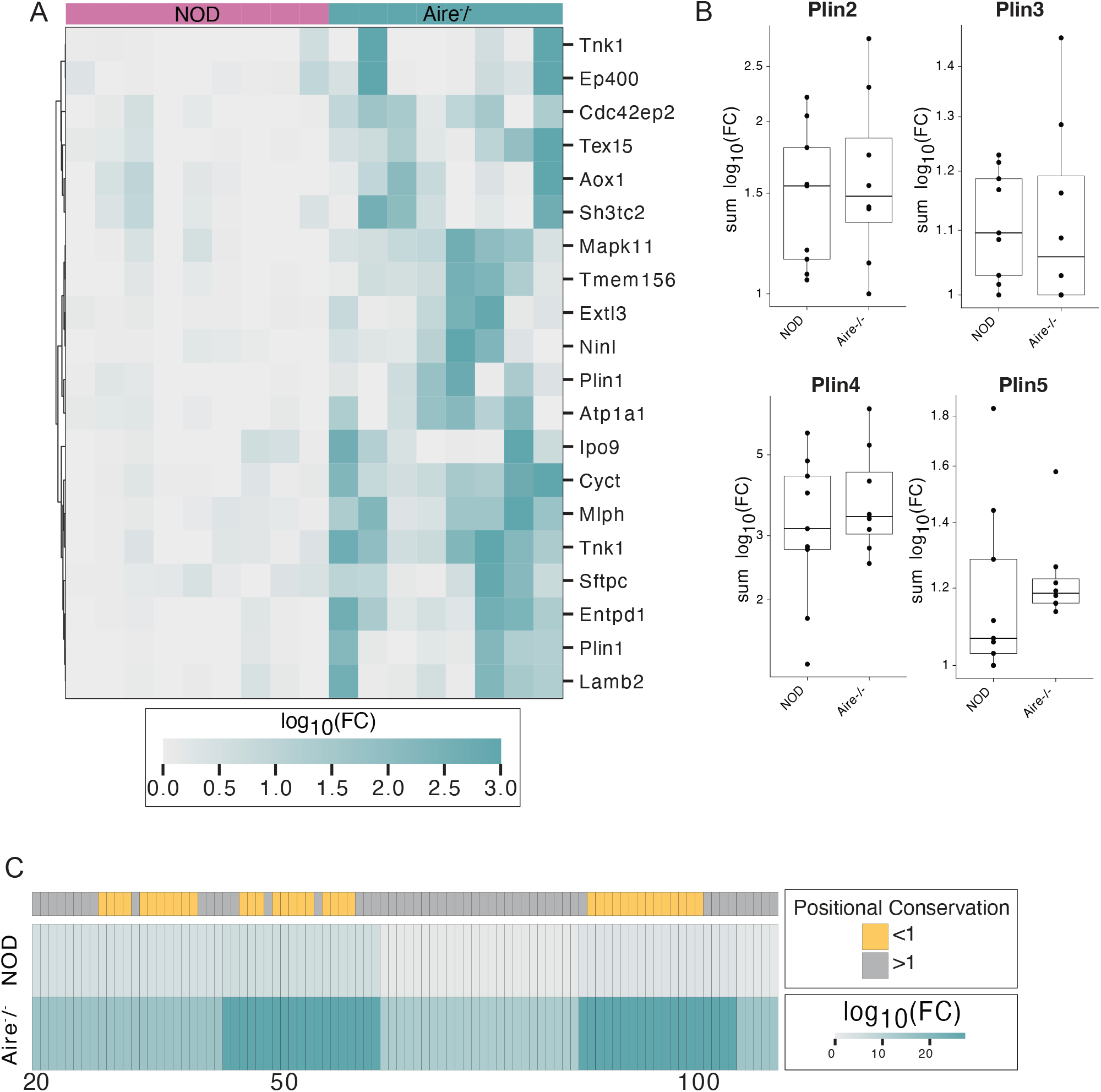
Autoreactive peptides in *Aire*^-/-^ mice. **A.** Heatmap of log_10_ fold change over background in top 20 peptides ranked by fold change in *Aire*^-/-^ versus NOD mice. **B.** Sum log_10_ fold change over MBM of peptides tiling across Plin2, Plin3, Plin4, and Plin5. **C.** Heatmap of Plin1 PAT domain positional sum log_10_ fold change over MBM in *Aire*^-/-^ or NOD mice annotated with positional conservation with other perilipin family proteins. Peptide enrichments were identified by PhIP-seq and annotated by their corresponding protein.

**Figure S6.**
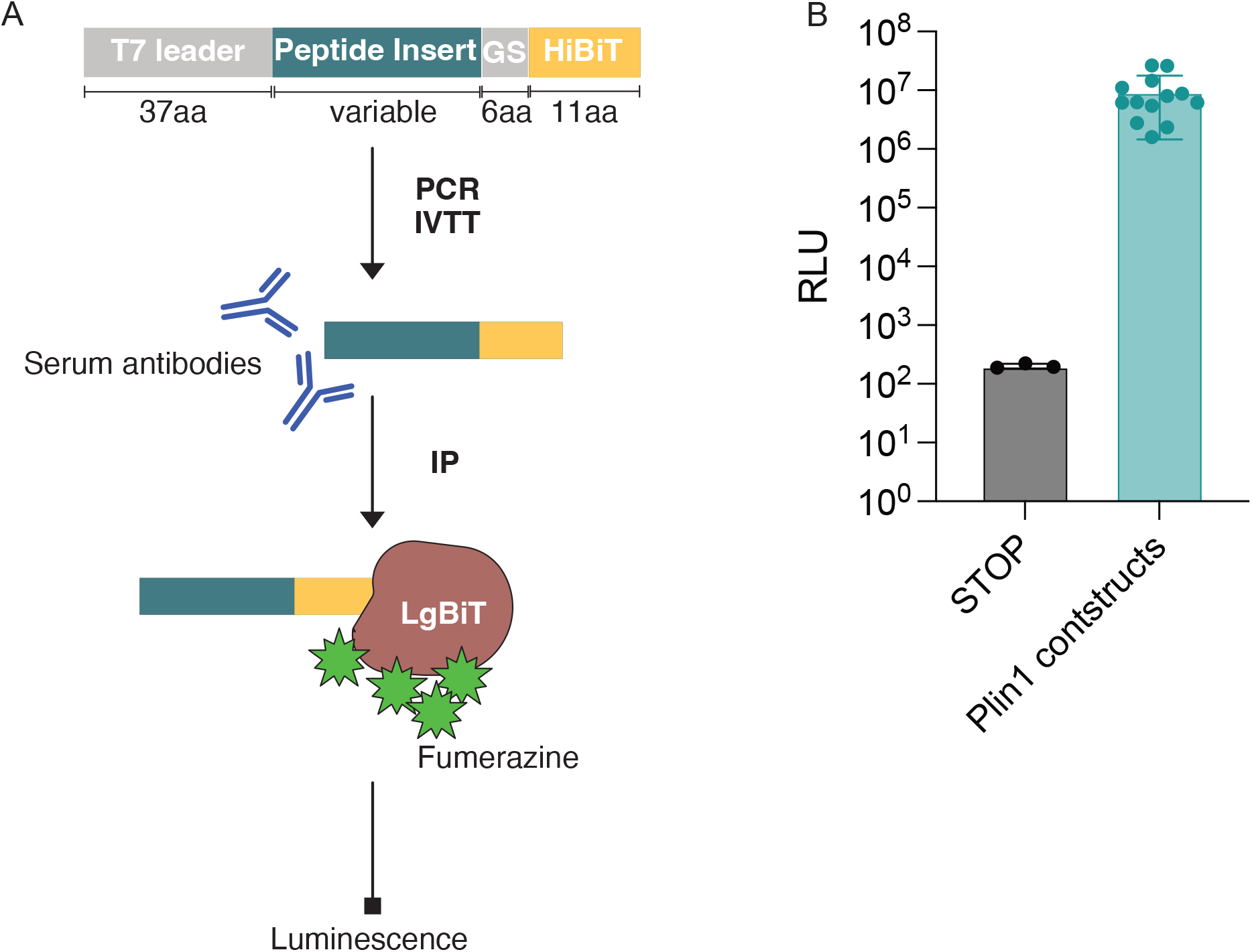
Method for split luciferase binding assay. **A.** Schematic of split luciferase binding assay (SLBA) protocol. HiBiT-tagged constructs are synthesized as DNA oligomers, amplified by PCR, and *in vitro* translated (IVTT). After immunoprecipitation (IP) with serum antibodies, peptide enrichment is quantified by adding LgBiT that complexes with HiBiT tag and generates luminescence given a fumerazine substrate. **B.** Relative luminescence units (RLU) of in-frame stop codon or Plin1 constructs.

**Figure S7.**
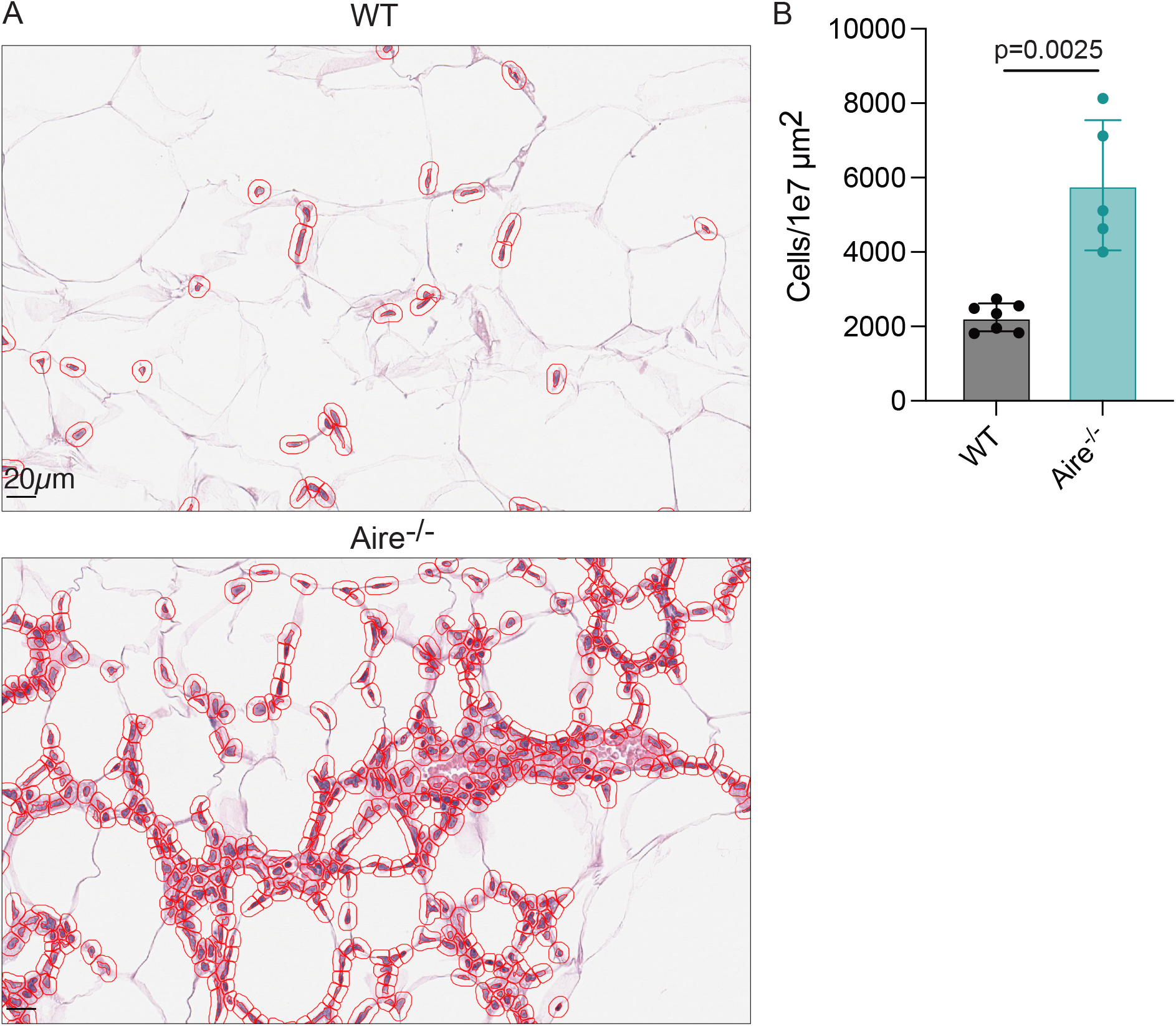
Quantification of cell-infiltrates in adipose tissue of *Aire*^-/-^ mice. **A.** Representative images of inguinal fat pads stained with H&E and infiltrating cell boundaries (red) identified by QuPath software in wildtype NOD or *Aire*^-/-^ NOD mice. Scale bar corresponds to 20µm. **B**. Infiltrating cells per 1e7µm^2^ area in wildtype NOD or *Aire*^-/-^ NOD mice. KS test for significance.

